# SpaGene: A Deep Adversarial Framework for Spatial Gene Imputation

**DOI:** 10.1101/2025.10.03.680242

**Authors:** Aishwarya Budhkar, Juhyung Ha, Qianqian Song, Jing Su, Xuhong Zhang

## Abstract

Integrating transcriptome-wide single-cell gene expression data with spatial context significantly enhances our understanding of tissue biology, cellular interactions, and disease progression. Although single-cell RNA sequencing (scRNA-seq) provides high-resolution gene expression data, it lacks crucial spatial context, whereas spatial transcriptomics techniques offer spatial resolution but are limited in the transcriptomic coverage. To address these limitations, integrating scRNA-seq and spatial transcriptomics data is essential. We introduce SpaGene, a novel deep learning framework designed to integrate scRNA-seq data and spatial transcriptomics data. SpaGene consists of two encoder-decoder pairs combined with two translators and two discriminators to effectively impute missing gene expressions within spatial transcriptomics datasets. We benchmarked SpaGene against existing state-of-the-art methods across diverse datasets. Across the datasets, SpaGene achieved an average 33% higher Pearson correlation coefficient (PCC), 21% higher Structural similarity index (SSIM), and 6.6% lower Root mean squared error (RMSE) compared to the existing approaches, highlighting its capability to reliably impute missing genes and provide comprehensive transcriptomics profiles. Application of our model to lung tumor tissue revealed immune cell enrichment at tumor boundaries, restricted myeloid cell trafficking in adjacent normal regions, and microenvironmental-driven pathways linked to immune neighborhoods. These results provide novel insight into immune exclusion and tumor-immune interactions that drive tumor progression, highlighting potential avenues for therapeutic development. Thus, SpaGene extends the power of spatial transcriptomics by delivering spatially resolved, enhanced transcriptome data that enable deeper biological understanding.

## INTRODUCTION

Recent advances in spatial transcriptomics (ST) techniques now make it possible to capture gene expression at the single-cell level while retaining spatial information about cells. For example, using positional barcodes, detected RNA transcripts can be mapped back to their tissue region to retain spatial information^1^. Understanding transcriptional profiles at the cellular level aids in several ways, such as recognition of the similarities and differences within cell populations, which helps elucidate cellular heterogeneity, identification of cell development pathways, study of rare cell populations such as tumor cells, etc^2, 3^. The availability of commercial platforms like the Vizgen MERSCOPE platform^4^ and the NanoString CosMX™ Spatial Molecular Imager (SMI) platform^5^ has enabled researchers and clinicians to access high-resolution ST data, facilitating the discovery of novel biological insights^3^. Spatial information, along with gene expression profiles, helps biologists understand complex cellular relationships and the resulting biological phenomenon^6^. For example, the NanoString platform enables spatial in situ detection of mRNA and proteins at the cellular and subcellular levels using formalin-fixed paraffin-embedded (FFPE) and fresh frozen (FF) tissue samples^5^. Another imaging-based technique, MERSCOPE, uses Multiplexed Error Robust Fluorescence In Situ Hybridization (MERFISH) to capture the spatial distribution of RNA molecules at the single-cell level. However, different ST techniques have their own limitations, such as less accurate capture of gene expression or limited spatial resolution. For example, imaging-based techniques such as MERFISH^7^ and osmFISH^8^ provide single-cell resolution with high accuracy. Still, they are limited to measuring gene expressions for hundreds to a few thousand genes. Sequencing-based techniques like Slide-seq^9^ and 10x Visium^10^ can detect thousands of genes, but at lower spatial resolution than single-cell techniques and with lower capture efficiency. Similarly, the NanoString CosMX™ SMI platform^5^ can profile thousands of genes, but the detection accuracy is low due to limitations of the technology.

Given that current ST technologies face notable limitations, there is a pressing need for computational strategies to enhance the quality and resolution of ST data. Prior to the development of the ST technology, single-cell RNA sequencing (scRNA-seq) emerged as a powerful tool for dissecting cellular heterogeneity and tracing cell lineages^2, 3^. However, while scRNA-seq (SC) offers detailed molecular profiles, it lacks spatial information, making it difficult to reconstruct the tissue architecture and cell-cell interactions within complex biological systems^1^. When combined with ST, SC serves as a valuable complementary modality, boosting the accuracy of spatially resolved transcriptomic analyses within individual tissue sections. Several computational methods have been proposed to integrate SC with ST for gene expression imputation to overcome ST’s inherent limitations. For example, SpaGE (Spatial Gene Enhancement)^11^ employs the PRECISE^12^ domain adaptation algorithm to align datasets. After alignment, the *k*-nearest neighbor algorithm^13^ (*k*-NN) is used to assign gene expression to spatially unmeasured locations using a weighted average of neighbors with positive cosine similarity. However, SpaGE faces limitations in handling complex datasets due to its reliance on *k*-NN and the PRECISE^12^ algorithm, which uses linear dimensionality reduction. gimVI (Gene imputation with Variational Inference)^14^ utilizes a deep generative model to integrate ST and SC datasets. The method learns a shared latent space of the input datasets and then uses posterior inference for gene imputation. gimVI can struggle to capture nuanced biological variations due to its limitations in latent space complexity and variability modeling. Tangram^15^ integrates ST and SC data by learning a probabilistic assignment of cells to spatial spots and uses the mapping to impute gene expression at cellular resolution. However, its effectiveness depends on shared gene expression patterns between datasets, which can limit performance in highly heterogeneous or novel biological contexts.

In this work, we introduce SpaGene, a novel deep learning method for predicting unmeasured genes in ST data by leveraging information from SC data. SpaGene utilizes an advanced encoder-decoder architecture supplemented by dedicated translator and discriminator modules to accurately impute missing gene expressions within ST data. First, the encoder-decoder modules project each dataset into a low-dimensional latent space and reconstruct it back to the original dimension, learning to capture the distinctive characteristics of each dataset while preserving biologically relevant variation. Then, translators learn mappings between latent spaces, learning their shared features and their relationships. Discriminator modules further refine these mappings by encouraging the translated features to resemble the target data distribution. By leveraging the learned features from both datasets, SpaGene accurately imputes expression for unmeasured genes in ST data, significantly enhancing the analytical power of ST. This enables more comprehensive downstream analysis, such as spatial cellular interaction, pathway enrichment analysis, and microenvironment profiling, leading to advancing our ability to extract biological insights from spatial data.

## RESULTS

### Overview of the SpaGene model

The primary function of the model is to enhance the limited ST data by expanding measured gene sets, significantly enriching the underlying biological information and, thus, facilitating novel biological insights through downstream analysis and interpretation. **Figure 1a** illustrates the overall objective of SpaGene. SpaGene leverages reference SC data to predict unmeasured genes in ST datasets, resulting in spatial profiles enriched with imputed genes. As shown in **Figure 1b**, the model comprises three components: encoder-decoder modules, translators, and discriminators. The model uses separate encoder-decoder networks for ST and SC datasets to capture and encode the unique dataset-specific features, mapping each dataset to compact, low-dimensional latent representations. This step is crucial to effectively capture essential characteristics of complex biological datasets and mitigate noise. The translators and discriminators then learn intricate non-linear mappings between the latent spaces of ST and SC datasets. Translators are trained to convert representations from one dataset to another, while discriminators guide the translators to generate realistic outputs. This adversarial framework encourages translated outputs that are biologically plausible and closely resemble the target data distributions. The discriminators enforce stringent quality control on translated representations, helping translators to iteratively refine their outputs towards increased biological relevance. **Figure 1c** further elaborates on the domain translation process, where ST and SC data are projected into their latent representations and translated across domains. This bidirectional translation using the learned low-dimensional latent space representation helps the model learn shared features between the two datasets. As depicted in **Figure 1d** during inference, ST data is projected into the learned latent space and translated into the SC domain. Finally, the gene expression is reconstructed, enhancing the ST data by expanding the measured gene expression. Thus, leveraging both shared and unique features of both datasets, SpaGene learns to translate ST data into the SC domain. Through adversarial training, SpaGene ensures accurate alignment and imputation.

**Figure 1.**
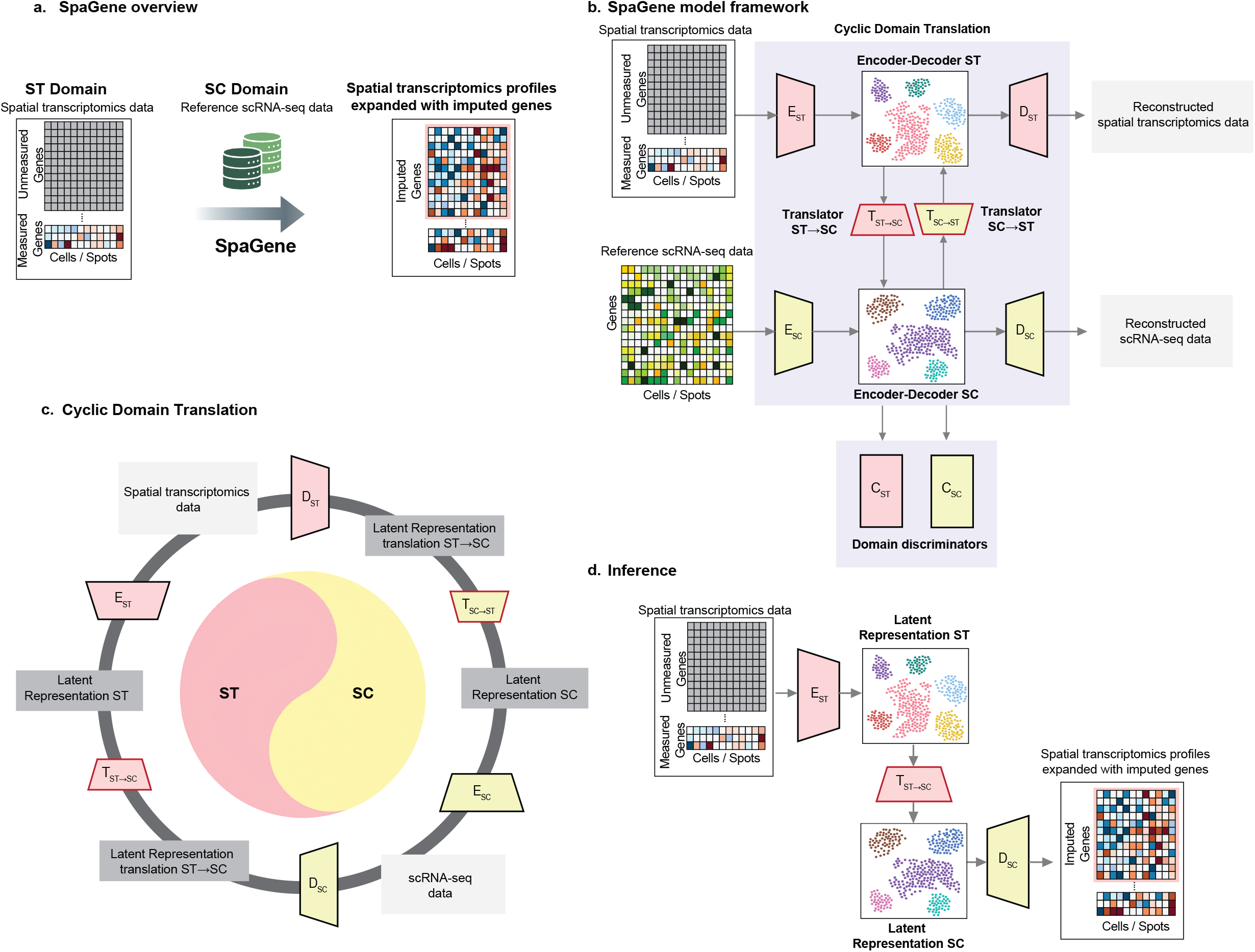
Overview of the SpaGene model. (a) SpaGene take ST data and reference SC data as input to impute unmeasured genes in ST data. (b) SpaGene consists of two encoder-decoder pairs for ST and SC data each to learn the unique features of the datasets, two translators to map the latent representations between datasets, and two discriminators that guide the translators to generate realistic, biologically meaningful outputs. (c) Data are first embedded into a low-dimensional latent representation, followed by translation to the target dataset and back to the source to learn the shared features across datasets. (d) Missing gene expression in ST is imputed by translating the latent representation to the SC domain and decoding it to reconstruct the gene expression.

### SpaGene outperforms existing methods across diverse datasets

To assess SpaGene’s ability in imputing missing gene expression, we benchmarked it against leading data imputation methods, gimVI^14^, SpaGE^11^, and Tangram^15^ across seven diverse dataset pairs: MERFISH^7^_Moffitt^7^, NanoString^5^_GSE^16^, osmFISH^8^_AllenSSp^17^, osmFISH_AllenVISp^18^, osmFISH_Zeisel^19^, seqFish^20^_AllenVISp, and STARmap^21^_AllenVISp detailed in the Methods section. Performance was evaluated using three metrics: Pearson correlation coefficient^22^ (PCC), structural similarity index^22^ (SSIM), and root mean square error^22^ (RMSE). Across all seven benchmarks, SpaGene consistently outperforms SpaGE, gimVI and Tangram by capturing non-linear relationships and more faithfully reconstructing smother, more accurate gene expression patterns.

**Figure 2a** shows the PCC results for each method across the seven datasets. SpaGene yields stronger agreement with ground-truth spatial measurements. For example, on the imaging-based MERFISH_Moffitt dataset pair, SpaGene achieved an average PCC of 0.3966, outperforming SpaGE (PCC = 0.3447), gimVI (PCC = 0.2418), and Tangram (PCC = 0.2727). Similarly, on the Nanostring_GSE dataset pair, SpaGene achieved an average PCC of 0.2456, 67.1% higher than SpaGE (PCC = 0.1469), 48.7% higher than gimVI (PCC = 0.1652), and 35.2% higher than Tangram (PCC = 0.1817). Thus, SpaGene produces stronger alignment between predicted and measured expression profiles of individual genes, reflecting its capacity to learn complex, non-linear mappings that faithfully reconstruct individual gene expression in spatial context. **Figure 2b** shows that SpaGene achieves higher SSIM scores than other methods across all seven datasets. This demonstrates that it produces a more precise recovery of spatial features, such as localized hotspots, than competing methods. For example, for the osmFISH_AllenVISp dataset pair, SpaGene achieved an average SSIM of 0.3821, better than SpaGE (average SSIM = 0.2268), gimVI (average SSIM = 0.3235), and Tangram (average SSIM = 0.2528). Similarly, for the osmFISH_Zeisel dataset pair, SpaGene achieved an average SSIM of 0.4127, 30.6% higher than SpaGE (SSIM = 0.3159), 37.9% higher than gimVI (SSIM = 0.2992), and 20.1% higher than Tangram (SSIM = 0.3434). These results show that SpaGene not only captures gene expression more accurately but also better preserves spatial architectures underlying tissue organization. **Figure 2c** presents RMSE values across all seven datasets. SpaGene consistently reduces the average discrepancy between predicted and measured gene expression intensities across all datasets. For example, for the seqFISH_AllenVISp dataset pair, SpaGene achieved an average RMSE of 1.1613, better than SpaGE (RMSE = 1.2435), gimVI (RMSE = 1.2570), and Tangram (RMSE = 1.2235). Similarly, for the STARmap_AllenVISp dataset pair, SpaGene achieved an average RMSE of 1.2496, lower than SpaGE (average RMSE = 1.3083), gimVI (average RMSE = 1.2763), and Tangram (average RMSE = 1.2773). Lower reconstruction error highlights SpaGene’s ability to minimize systematic and random deviations, yielding intensity predictions that closely align with observed data.

**Figure 2.**
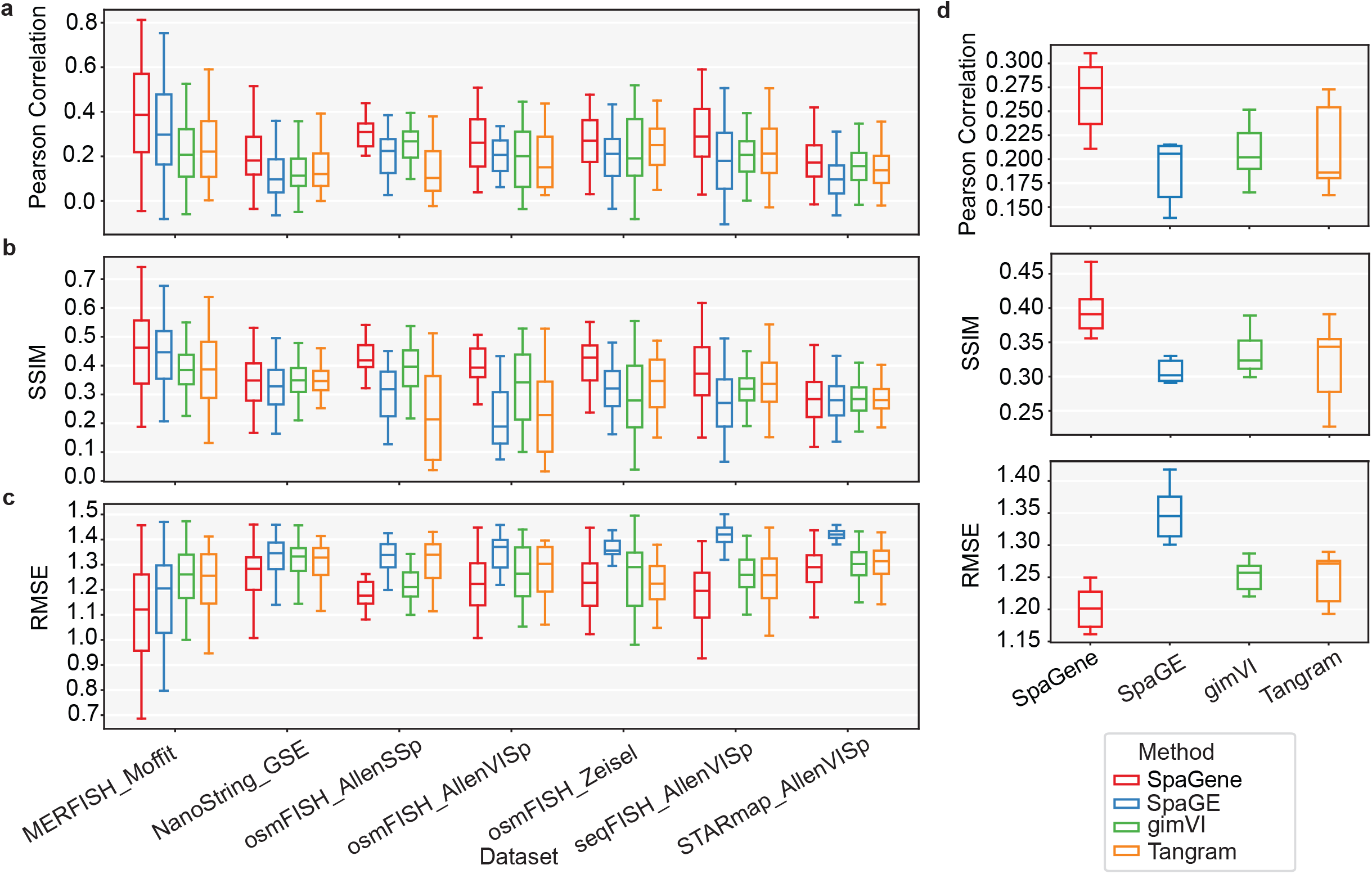
Performance evaluation across datasets. (a) Box plot of PCC scores across seven datasets for each method (b) Box plot of SSIM scores across seven datasets for each method (c) Box plot of RMSE scores across seven datasets for each method (d) Box plot of average PCC, SSIM, and RMSE scores for the seven datasets for each method

Across all seven datasets, SpaGene achieved an average PCC of 0.2782, 36% higher than SpaGE (PCC = 0.2045), 34% higher than gimVI (PCC = 0.2076), and 30.7% higher than Tangram (PCC = 0.2129), average SSIM of 0.3902, 23.6% higher than SpaGE (SSIM = 0.3158), 16.8% higher than gimVI (SSIM = 0.3342), and 22.7% higher than Tangram (SSIM = 0.3180), and a lower RMSE score of 1.1898, 10.1% lower compared to SpaGE (RMSE = 1.3238), 4.97% lower than gimVI (RMSE = 1.2520), and 4.6% lower than Tangram (RMSE = 1.2472) as shown in **Figure 2d**. Collectively, these results demonstrate SpaGene’s robust performance across diverse datasets including large-scale imaging-based datasets like MERFISH profiling thousands of cells with large gene panels to targeted osmFISH assays where spatial measurements are limited to a small gene panel. Thus, the model’s architecture is suitable to handle both high and low-dimensional data. This adaptability shows that SpaGene’s underlying mechanisms are not dependent on technology or gene counts but exploit underlying patterns in spatial and single-cell data to generate reliable imputations. Overall, the SpaGene framework produces more accurate, visually coherent, and quantitatively reliable gene imputations that reflect true underlying biological mechanisms.

### SpaGene demonstrates superior performance on the NanoString Lung9 rep1 dataset

We applied SpaGene to integrate the NanoString Lung9 rep1 dataset with reference SC data to predict unmeasured spatial gene expression patterns, enriching the ST data. To rigorously evaluate the predictive performance of SpaGene, we conducted a five-fold cross-validation, detailed in the Methods section. In each fold, a subset of genes was held out, and the remaining genes were used to impute the spatial expression of the omitted genes. We compared our model’s performance with SpaGE, gimVI, and Tangram using PCC between measured spatial gene expression and predicted values. **Figure 3a** shows the PCC results for the NanoString Lung9 rep1 tissue sample, highlighting the superiority of SpaGene. SpaGene achieved an average PCC of 0.2456, significantly higher than SpaGE (average PCC = 0.1470), gimVI (average PCC = 0.1652), and Tangram (average PCC = 0.1817). We further conducted gene-level comparisons to provide detailed insights into the method performance. **Figures 3b-d** show scatter plots comparing gene-wise PCC values between SpaGene and each competing method. We observed that the majority of the data points lie above the *y = x* line, demonstrating that SpaGene performs better than the competitors.

**Figure 3.**
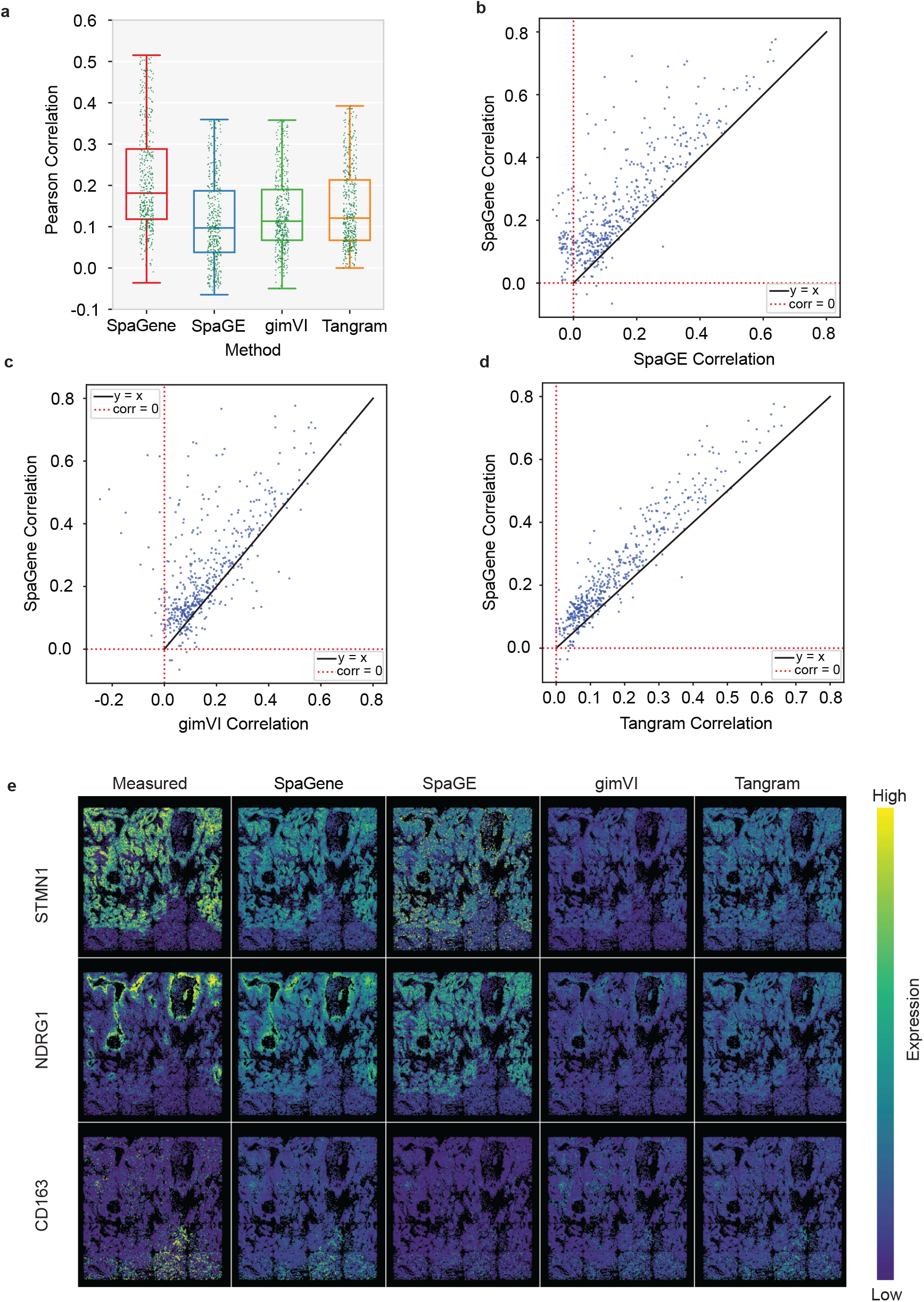
Performance evaluation on the NanoString Lung9 rep1 sample. (a) Box plot of PCC for each method (b) Scatter plot of PCC values for each imputed gene between SpaGene versus SpaGE (c) Scatter plot of PCC values for each imputed gene between SpaGene versus gimVI (d) Scatter plot of PCC values for each imputed gene between SpaGene versus Tangram (e) Spatial patterns of measured and imputed genes *STMN1, NDRG1*, and *CD163* for each method

Furthermore, we visually assessed the spatial expression patterns of selected imputed genes with complex spatial patterns. **Figure 3e** illustrates measured and imputed spatial patterns of *STMN1, NDRG1*, and *CD163* genes in the NanoString Lung9 rep1 sample. Compared with SpaGE, gimVI, and Tangram, SpaGene consistently produced more spatially coherent patterns that closely matched the measured patterns. For example, SpaGene accurately captured the intricate spatial heterogeneity of *NDRG1*, and *CD163* clearly outperforming the competitors. SpaGE produced a noisy pattern for *STMN1* and failed to capture the complex spatial distribution for *NDRG1* and *CD163*. gimVI overly smoothed the expression pattern for *STMN1*, and Tangram failed to reproduce the spatial expression pattern for *NDRG1*. The robustness of SpaGene is evident in accurately reconstructing distinct spatial expression patterns, which suggests reliable performance.

### SpaGene demonstrates superior performance on the STARmap dataset

Similarly, we evaluated the performance of SpaGene on the STARmap_AllenVISp dataset in predicting unmeasured spatial gene expression patterns. To assess the predictive accuracy, we used five-fold cross-validation and quantified the performance using the PCC metric. **Figure 4a** summarizes the PCC results for the STARmap data, where SpaGene achieved an average PCC of 0.2108, outperforming SpaGE (PCC = 0.1386), gimVI (PCC = 0.1798), and Tangram (PCC = 0.1786). This marked improvement highlights the strong predictive power of SpaGene. To examine gene-level performance, **Figures 4b-4d** present scatter plots comparing SpaGene’s PCC values with those of the other methods. Across all competitors, more data points lie above *y = x* line, which demonstrates that SpaGene often yields higher PCCs for a large proportion of genes.

**Figure 4.**
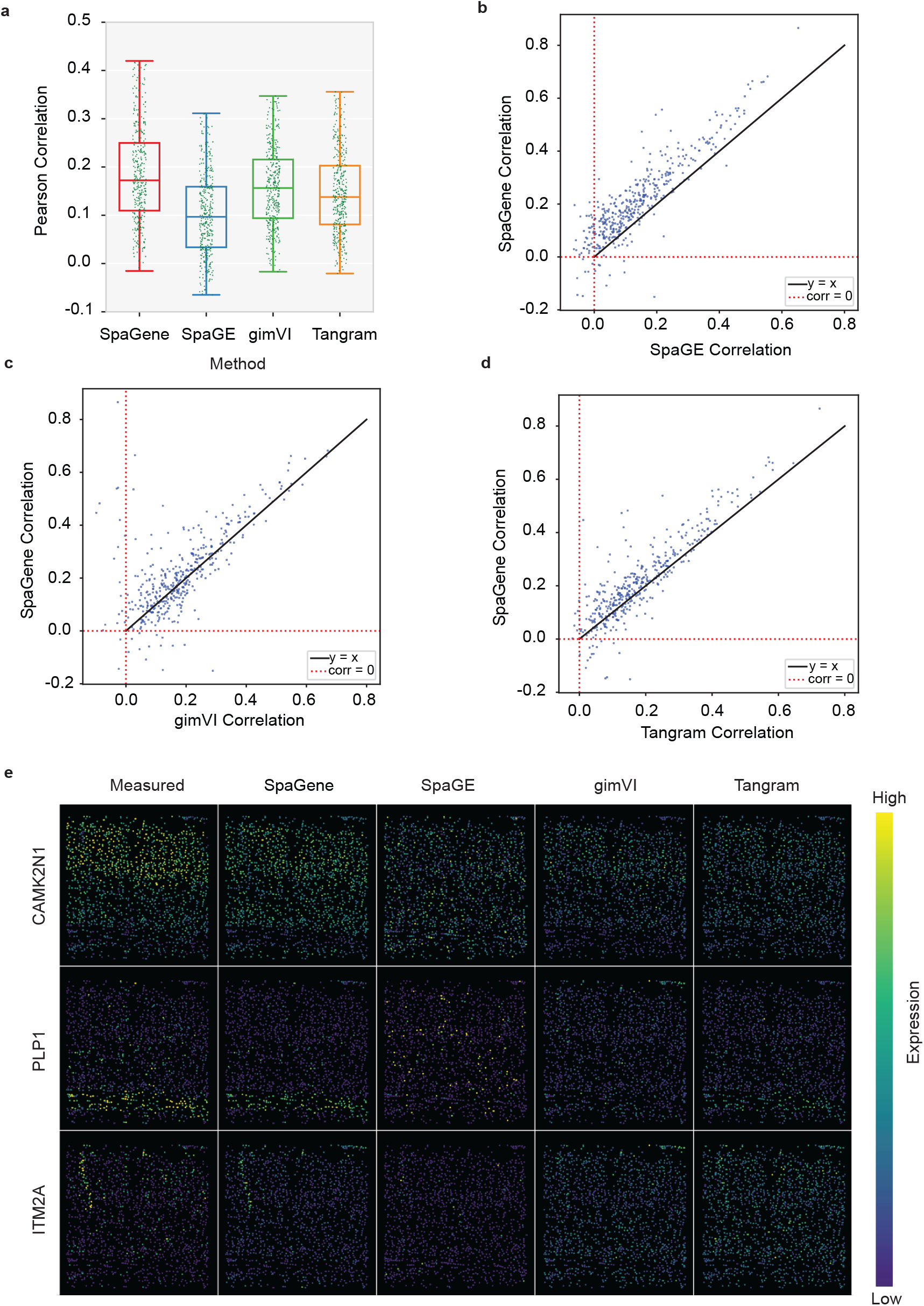
Performance evaluation on the STARmap data. (a) Box plot of PCC is shown for each method (b) Scatter plot of PCC values for each imputed gene between SpaGene versus SpaGE (c) Scatter plot of PCC values for each imputed gene between SpaGene versus gimVI (d) Scatter plot of PCC values for each imputed gene between SpaGene versus Tangram (e) Spatial patterns of measured and imputed *CAMK2N1, PLP1*, and *ITM2A* genes for each method

In addition to quantitative performance, we evaluated the spatial fidelity of imputed gene expression patterns. **Figure 4e** displays spatial patterns of three representative imputed genes, *CAMK2N1, PLP1*, and *ITM2A*, by showing the ground truth STARmap measurements alongside predicted expression patterns for SpaGene, SpaGE, gimVI, and Tangram. We observed that gene expression imputed by SpaGene more closely replicates the measured spatial distributions compared to the competitors. For example, for *CAMK2N1*, SpaGene captures spatial heterogeneity of the expression pattern better than gimVI, which performs reasonably well but suppresses some expression patterns. SpaGE and Tangram lose the spatial heterogeneity of the expression pattern in some regions. For *PLP1*, SpaGene closely matches the measured spatial pattern, Tangram captures the pattern reasonably well, whereas gimVI and SpaGE fail to do that. For *ITM2A*, SpaGene shows a consistent spatial pattern with measured expression, while others cannot accurately reproduce the pattern. Together, these visual and quantitative results establish that SpaGene delivers more accurate and biologically realistic spatial imputation compared to existing methods.

### SpaGene leverages imputed genes to uncover novel biological insights

We investigated how the local immune microenvironment influences tumor cell states in the NanoString Lung9 rep1 sample using imputed gene expression profiles with spatial neighborhood-based analysis. First, we defined spatial niches using Seurat v5^23^ based on each cell’s local neighborhood composition using k-nearest neighbors. Five major niches in the microenvironment were identified: tumor, myeloid, fibroblast, neutrophil, and lymphocyte. **Figure 5a** displays the spatial plot of cells with cell types and identified spatial niches that capture functionally distinct regions within the tissue.

**Figure 5.**
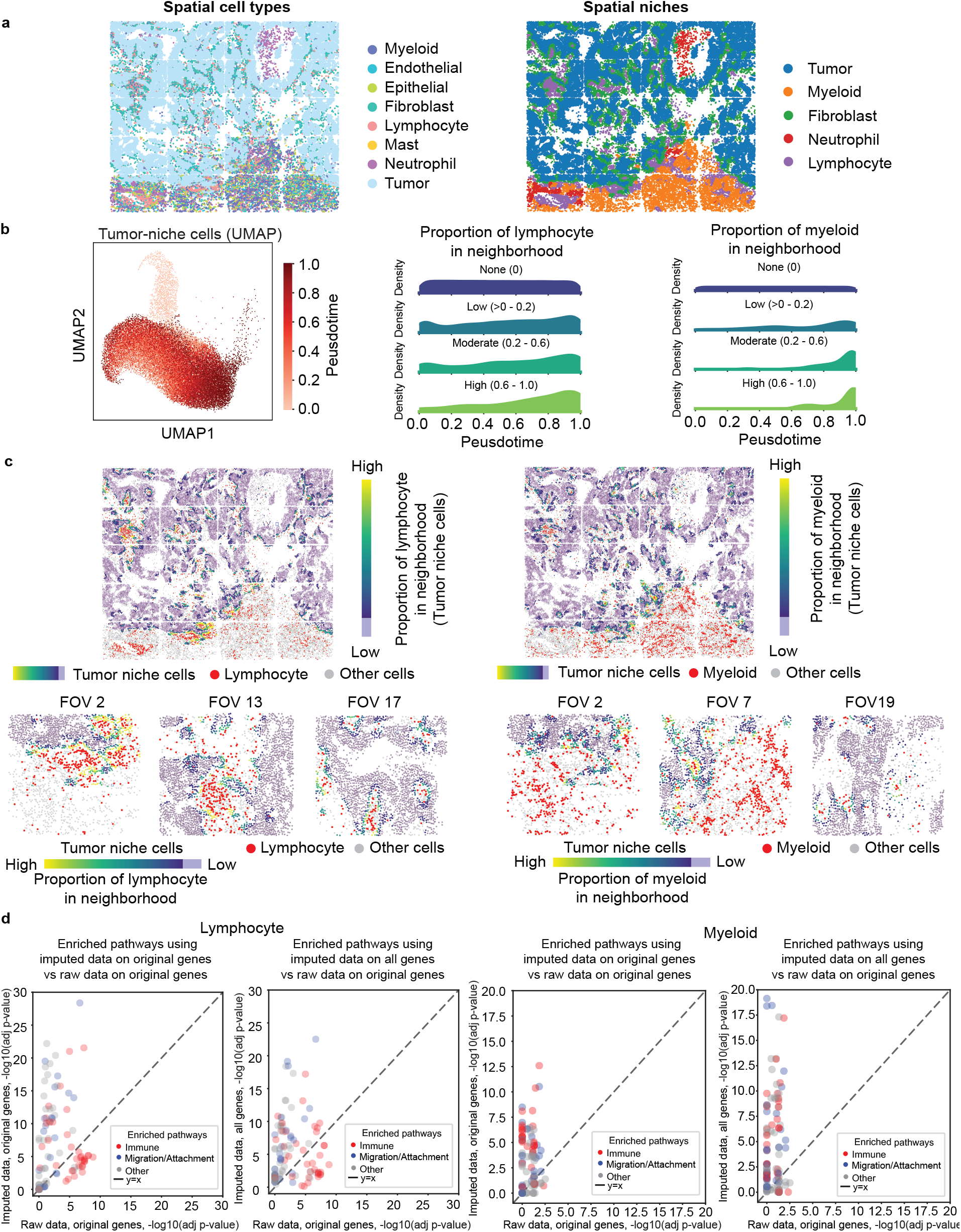
Downstream analysis using imputed data. (a) Spatial niche analysis. Spatial plot of cells colored by cell type, Spatial plot of cells colored by spatial niche: tumor, myeloid, fibroblast, neutrophil, and lymphocyte. (b) Pseudotime and immune neighbor association among tumor niche cells. UMAP projections for tumor niche cells colored by the inferred pseudotime (ptime), Ridge density plots showing distribution of tumor niche cells along pseudotime, grouped based on their lymphocyte and myeloid cell neighbor proportion (none (0), low (>0-0.2), moderate (0.2-0.6), high (0.6-1.0)). (c) Immune neighborhood for tumor niche cells. Spatial plot of cells colored by proportion of lymphocyte in neighborhood (left) and proportion of myeloid in neighborhood (right) for tumor niche cells with immune cells of interest highlighted in red and other cells in gray. Representative fields of view (FOVs) for lymphocyte (FOVs 2, 13, 17) and myeloid (FOVs 2, 7, 19) neighbor proportions for tumor niche cells colored from high (yellow) to low (purple), immune cells in red, and other cells in grey. (d) Pathway enrichment comparisons. Scatter plots of pathway enrichment significance (–log_10_ adj p-value) detected using imputed versus raw expression data for lymphocyte (left two panels) and myeloid (right two panels): imputed original (only measured genes in raw data) versus raw (first column) and imputed all (both measured genes in raw data and newly imputed genes) versus raw (second column). Points in scatter plots denote pathways colored by annotation – Immune (red), Migration/Attachment (blue), Other (gray), and the dashed line indicates the *y=x* line.

To study tumor cell progression, we applied SpaTrack^24^ to infer pseudo-temporal trajectories within tumor niche cells. **Figure 5b** presents UMAP embeddings of tumor niche cells colored by inferred pseudotime and ridge density plots depicting distributions for these cells along pseudotime, grouped by their lymphocyte and myeloid cell neighbor proportion (none (0), low (>0-0.2), moderate (0.2-0.6), high (0.6-1.0)). The ridge density plots reveal that tumor niche cells at later pseudotime are enriched in immune-rich neighborhoods. These patterns suggest that immune infiltration is associated with tumor state transitions^25, 26^. To spatially validate these patterns, we visualize the spatial plot of tumor niche cells in **Figure 5c**. Tumor niche cells are colored by their lymphocyte and myeloid neighbor proportion, with immune cells of interest highlighted in red and all other cells shown in gray. Lymphocyte-rich neighborhoods are seen to be localized near tumor boundaries, while myeloid-rich neighborhoods are predominantly found in supporting tissue around the tumor. Representative fields of view (FOVs) illustrate immune enrichment at the tumor-normal interface and show restricted deeper immune entry into tumor regions, with myeloid cells accumulating in the normal tissue around the tumor.

Next, we investigate molecular pathways associated with immune infiltration. Local immune cell abundance can drive transcriptional changes in neighboring cells^27, 28^. Therefore, we used the proportion of local immune neighbors for tumor niche cells to select genes for pathway analysis to capture both cell-specific and microenvironmentally regulated pathways that might not be captured by cell-type information alone. Specifically, we computed PCC between gene expression and local immune neighbor proportions (lymphocyte and myeloid) for each tumor niche cell. Genes with moderate, biologically meaningful correlations were selected independently from three data subsets: raw gene expressions, imputed gene expressions with genes present in raw data, and all imputed gene expressions. Using genes selected from each data subset, we performed pathway enrichment using Reactome and Gene Ontology (GO) gene sets from the MSigDB database collection^29, 30^. We identified significant pathways and combined the top 20 enriched pathways for each data subset, and manually annotated them as Immune, Cell Migration/Attachment related, or Other. **Figure 5d** shows scatter plots comparing enrichment significance (–log_10_ adj p-value) between raw and imputed data. We observed that using imputed expression increased the sensitivity for the detection of Immune and Cell Migration/Attachment related pathways compared with raw expression. Moreover, inclusion of newly imputed genes increased the significance of detected pathways compared with using imputed expression only for genes already measured in raw data. This demonstrates that imputed expression uncovers microenvironment-driven biological processes missed in raw data and improves the pathway detection sensitivity.

These results demonstrate how imputation enhances ST data, offering insights into tumor trajectory mapping, spatial immune tumor organization, and improved detection of biological pathways, in turn advancing our understanding of the complex tumor microenvironment.

## DISCUSSION

Spatial single-cell transcriptomics data plays an important role in understanding complex tissue structures and functions by revealing spatially resolved gene expression at single-cell resolution. However, only a limited subset of genes is typically captured, constraining comprehensive biological interpretation. Accurate gene imputation using single-cell reference data can address this limitation and aid in advancing spatial biology, enabling novel insights into cellular heterogeneity, interactions, and underlying tissue mechanisms. SpaGene addresses this critical need, serving as an advanced computational approach to enhance ST datasets, empowering researchers to gain novel biological insights.

The primary objective of SpaGene is to enhance the ST data by expanding gene coverage beyond the assayed set through effective integration with reference SC data. SpaGene imputes expression for unmeasured genes in ST data, significantly enhancing the biological signal and thereby facilitating downstream analysis to discover novel insights. The framework consists of two encoder-decoder networks, two translators, and two discriminators. Separate encoders learn robust latent space representations from ST and SC data. Two translators are trained to translate data between domains. To enhance ST data, its latent representation is translated into the SC domain by a dedicated translator module. The translated representation is then used to reconstruct comprehensive gene expression profiles using the SC decoder module. SpaGene’s superior performance stems from its adversarial framework that effectively captures shared information by learning non-linear mappings between source and target domains. By learning and leveraging complex data distributions, SpaGene improves the accuracy and reliability of gene imputation in spatial datasets, yielding comprehensive biologically meaningful transcriptome profiles. A key advantage of SpaGene is its ability to effectively capture complex biological signals and model the non-linear characteristics of omics datasets. This capacity is particularly beneficial for understanding spatial heterogeneity, intercellular communication, and functional cellular states. Thus, SpaGene stands out as a valuable tool for elucidating biological complexity at the spatial level.

Biologically, SpaGene provides a more comprehensive view of the tumor immune microenvironment using the expanded spatial transcriptomic profile, revealing cellular interactions and biological pathways underrepresented in raw data. In the NanoString Lung9 rep1 sample, the enriched transcriptome enabled detection of spatial patterns such as lymphocytes clustering at tumor boundaries and myeloid cells blocked at the boundary between tumor and normal region. The expanded gene coverage highlighted associations between the immune neighborhood and tumor cell state transitions, offering insights into how immune pressure influences tumor evolution. Moreover, the model improves detection of microenvironment-driven pathways, particularly immune and cell migration/ attachment pathways, relative to the raw data. These findings indicate that SpaGene utilizes the expanded transcriptome to provide insights into tumor evolution and to improve the identification of biologically relevant pathways that can be used to develop future therapeutic strategies.

While SpaGene has demonstrated superior performance in integrating ST and SC datasets, it holds potential for future research and development. One promising area of improvement involves the integration of additional data modalities such as imaging, proteomics, epigenomics, or metabolomics. Incorporating such multi-modal datasets would allow for a comprehensive understanding of tissue morphology and underlying biological processes. Such integration would enable a more holistic understanding of biological systems and potentially aid in the discovery of novel pathways and cellular interactions. Another essential avenue for future development is enhancing model interpretability through advanced explainability techniques^31^. Implementing attention frameworks^32^, integrated gradients^33^, layer-wise relevance propagation^34^, and related approaches could increase trust in model outputs. Greater transparency into how predictions are made could lead to the discovery of novel biomarkers, deeper insights into cellular mechanisms, and aid researchers in discovering hidden biological insights^31^. This will help for more effective translation of computational findings into biological knowledge. Overall, SpaGene is an advanced tool for spatial transcriptomics, significantly improving gene imputation and integration capabilities to drive forward research in spatial biology.

## MATERIALS AND METHODS

### Data processing

Following ST and SC dataset pairs are used for performance evaluation: MERFISH_Moffitt, NanoString_GSE, osmFISH_AllenSSp, osmFISH_AllenVISp, osmFISH_Zeisel, seqFish_AllenVISp, and STARmap_AllenVISp. For both ST and SC datasets, genes with density less than 0.05 and cells with density less than 0.1 were filtered out to reduce noise and computational load. Next, 2000 highly variable genes are selected using Scanpy^35^ v3. To reduce the influence of extreme values, outliers exceeding two standard deviations above the mean for each feature were clipped to the mean plus two standard deviations. Finally, we applied square root normalization by computing the square root of each expression value in the data to reduce data skewness.

### The SpaGene model

Our model consists of two Encoder-Decoder pairs, two Translators, and two Discriminators. Each Encoder-Decoder pair follows the AutoEncoder^36^ framework, which uses an encoder function to reduce the original data features into a low-dimensional feature space, and a decoder function to reconstruct the data from that low-dimensional feature space. The encoder is a multi-layer fully connected neural network with ReLU^37^ non-linear activations that compresses the given gene expression into a latent space representation. The decoder is a neural network that reconstructs the original gene expression from the latent space representation through successive nonlinear transformations. Translators facilitate domain adaptation by translating the latent space representation between the ST and SC domains. Discriminators act as binary classifiers for distinguishing whether a latent space representation originates from the source domain or was generated by a translator. The trained discriminator provides adversarial loss^38^ that guides the translators to generate more realistic outputs.

### Encoder-Decoder

SpaGene consists of two Encoder-Decoder modules, one for the ST and one for the SC domain. Each module comprises an encoder *E* that maps an input expression vector *x* to a low-dimensional latent representation and *z* decoder *D* that maps *z* back to a reconstruction 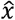. When applied to the ST and SC domains, the encoders produce *z*_*ST*_ = *E*_*ST*_ (*x*_*ST*_) and *z*_*SC*_ = *E*_*SC*_ (*x*_*SC*_), and the decoders reconstruct 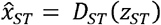 and 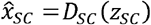 . To train the Encoder-Decoder, reconstruction error is minimized as follows:

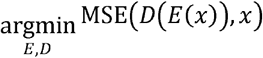

where MSE (x,y)stands for the mean-squared-error loss.

Both domain-specific modules are trained independently to minimize reconstruction error. Once the Encoder-Decoders are trained, their weights are frozen.

### Translator

The model comprises two directional translators: *T*_*ST*→*SC*_ that maps ST latent space *z*_*ST*_ to SC latent space *z*_*ST*→*SC*_ and *T*_*SC*→*ST*_ that maps SC latent space *z*_*SC*_ to ST latent space *z*_*SC*→*ST*_. Here,

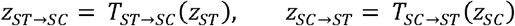

The objective of these translators is to generate realistic latent representations in the translated domains.

### Discriminator

We employ two discriminators: *C*_*ST*_ that classifies whether a latent representation is from real ST data or generated using the translator *T*_*ST*→*SC*_ and *C*_*SC*_ that distinguishes real SC embeddings from those produced by the translator *T*_*ST*→*SC*_ . Discriminators provide feedback that guides translators to generate more realistic representations through adversarial learning.

### Loss function

Multiple loss functions are employed, including cycle loss^39^, identity (ID) loss^40^, CORAL (CORrelation ALignment) loss^41^, MMD loss^42^, and GAN loss^40^.

Cycle loss: it ensures that translating the latent space to the other domain and back to the original domain preserves the original gene expression. Mathematically, the MSE between the original gene expression *x*_*ST*_ and the cycled gene expression 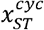 is minimized as follows:

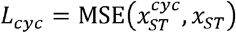

where MSE (x,y) is the mean-squared-error loss function. Thus, the cycle loss is defined as:

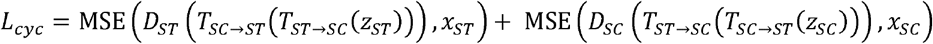

For identity (ID) loss, Pearson correlation between the original gene expression and the translated gene expression is used. Since the dimensions of the translated gene expression and original gene expression are different, backpropagation is only applied for shared genes that exist in both domains. By minimizing this loss, the translator learns to generate gene expressions that align closely with the target domain values. Thus, in our model, the ID loss is defined as:

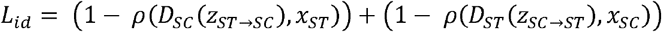

where *ρ* is the Pearson correlation computed over the shared gene set.

CORAL loss: it aligns the covariance between the ST and SC domains. The objective is to improve domain alignment in the latent space. CORAL loss minimizes the squared Frobenius norm of the difference between the covariance matrices of the two domains, as follows:

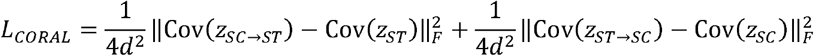

where ∥.∥_*F*_ is the Frobenius norm, Cov (·) denotes covariance, *d* and represents the number of features in the matrices to be aligned.

MMD (Maximum mean discrepancy) loss: this loss measures the distributional distance between latent space representations of ST and SC domains. By minimizing the MMD loss, the translator aligns the distributions of two domains in latent space to reduce domain shifts. The MMD loss for two distributions *P,Q* is given by:

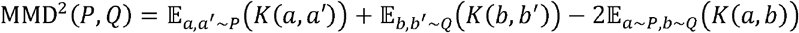

Where 𝔼 _*a,a*′ ∼*P*_ (*K*(*a,a*′)) is the expectation of the kernel function between two independent draws *a,a*′ from *P*, 𝔼 _*b,b*′ ∼*Q*_ (*K*(*b,b*′)) is the expectation of kernel function between two independent draws *b,b*′ from *Q*, 𝔼 _*a*∼*P*∼,*b*∼*Q*_(*K*(*a,b*)) is the expectation of kernel function between independent draws from *P,Q*, and *K* is Gaussian kernel. Thus, the MMD loss is defined as:

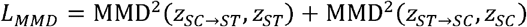

GAN loss: it introduces an adversarial objective where domain-specific discriminators are trained to classify whether the latent space representations originate from the source domain or are generated by the translator.

Each translator acts as the generator, mapping representations from one domain to the other with the goal of producing latent representations that are indistinguishable from real ones.

The translator is trained to fool the discriminator by generating representations *x*_*fake*_ that are indistinguishable from latent representations originating from the source domain *x*_*real*_ with the loss defined as:

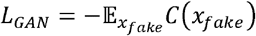

Specifically, for two translators the GAN loss is given as:

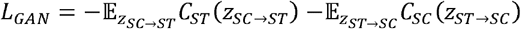

The discriminators are trained using hinge loss, enforcing a margin separator to distinguish between representations from the source domain and those generated by a translator as follows:

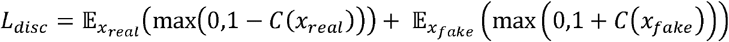

Thus, for our model, the discriminator loss is defined as:

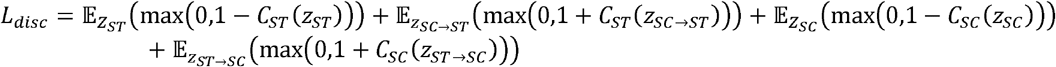

The translator loss combines ID loss, cycle loss, CORAL loss, MMD loss, and GAN loss as follows:

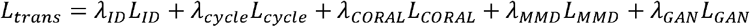

where*λ*_*ID*_,*λ*_*cycle*_,*λ*_*CORAL*_,*λ*_*MMD*_, *λ*_*GAN*_ are chosen using hyperparameter tuning.

### Model Inference

Gene expression from the ST domain *x*_*ST*_ is passed through the trained encoder *E*_*ST*_to generate the latent space representation *z*_*ST*_ ·*z*_*ST*_. is then translated into the SC domain *z*_*ST*→*SC*_ using the trained translator *T*_*ST*→*SC*_. Finally, the translated latent space is passed through the trained decoder of the SC domain *D*_*SC*_ to generate the translated gene expression from the ST to SC domain 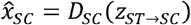.

### Experimental setup

The training is divided into two stages: In the first stage, the Encoder-Decoder networks are trained to efficiently encode and reconstruct gene expressions, ensuring faithful compression and reconstruction of input data. In the second stage, translators are trained to translate the latent representations from ST to SC. Multiple loss functions are utilized to ensure accurate and smooth domain translation including cycle loss, identity (ID) loss, CORAL loss, MMD loss, and GAN loss.

For cross-validated inference, the model is trained to predict a subset of genes held-out in each fold for evaluation. This is done by first encoding ST gene expression into a low-dimensional latent representation, translating that latent space to the SC domain, and finally decoding it to reconstruct full gene expression including genes not used during training in that fold. Through this process, the SpaGene model achieves its goal: translating gene expression data in the ST domain into the SC domain, thus enhancing the ST data and enabling more meaningful downstream analysis.

We evaluate SpaGene on seven ST and SC dataset pairs. For each fold in cross-validation, the model is trained on a subset of genes and evaluated on the held-out gene set. During each fold, the inference to translate held-out gene expression from ST to SC is done after training all components of the model including the two encoder-decoder pairs, two translators and two discriminators.

For each fold, the model is trained on four-fifths of the shared genes and tested on the remaining fifth, and this process is repeated five times. Each dataset is partitioned based on the set of shared genes between the ST and SC domains (MERFISH Moffitt dataset pair: 141, NanoString GSE dataset pair: 488, osmFISH AllenSSp dataset pair: 26, osmFISH AllenVISp dataset pair: 26, osmFISH Zeisel dataset pair: 32, seqFISH AllenVISp dataset pair: 411, STARmap AllenVISp dataset pair: 242). In each round of cross-validation, four gene folds are used for integration while the remaining fold is used for evaluation. Model performance was evaluated by comparing the measured and predicted gene expression profiles on the held-out genes.

### Evaluation metrics

Given *N* the number of cells and *G* the number of genes in the ST dataset, the measured expression for gene *g* as *x*_*g*_, and the predicted expression for gene *g* as 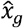, we used the following metrics to assess the performance of our method:

PCC: The Pearson correlation coefficient^22^ between each gene *g* in predicted expression by each method and the ground truth expression of ST is computed as follows:

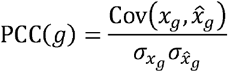

where 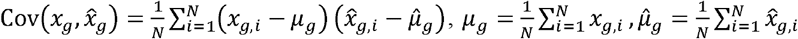, and 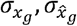 are their standard deviations.

SSIM (Structural similarity index): The structural similarity index^22^ measures the similarity between measured and predicted gene expression and is computed as follows:

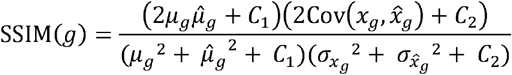

RMSE (Root mean square error): The root mean square error^22^ measures the error between measured and predicted gene expression and is computed as follows:

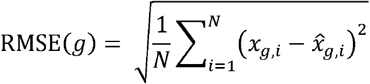

The average of PCC, RMSE and SSIM across held-out genes is reported for performance evaluation. Higher PCC and SSIM and lower RMSE indicate better prediction accuracy.

## DATA AVAILABILITY

GSE dataset can be download from https://www.ncbi.nlm.nih.gov/geo/query/acc.cgi?acc=GSE131907. NanoString CosMx SMI dataset can be download from https://nanostring.com/products/cosmx-spatial-molecular-imager/ffpe-dataset/, MERFISH Moffitt, osmFISH AllenSSp, osmFISH AllenVISp, osmFISH Zeisel, seqFISH AllenVISp, STARmap AllenVISp datasets can be download from the public repository https://zenodo.org/records/3967291.

## CODE AVAILABILITY

The SpaGene method is provided as an open-source Python package on GitHub: https://github.com/asbudhkar/SpaGene.

